# What does mitogenomics tell us about the evolutionary history of the *Drosophila buzzatii* cluster (*repleta* group)

**DOI:** 10.1101/712232

**Authors:** Nicolás N. Moreyra, Julián Mensch, Juan Hurtado, Francisca Almeida, Cecilia Laprida, Esteban Hasson

**Author notes:** Corresponding author (NNM); (EH).

## Abstract

The *Drosophila repleta* group is an array of more than 100 cactophilic species endemic to the “New World”. The acquisition of the ability to utilize decaying cactus tissues as breeding and feeding sites is a key aspect that allowed the successful diversification of the *repleta* group in the American deserts. Within this group, the *Drosophila buzzatii* cluster is a South American clade of seven cactophilic closely related species in different stages of divergence, a feature that makes it a valuable model system for evolutionary research. However, even though substantial effort has been devoted to elucidating the phylogenetic relationships among members of the *D. buzzatii* cluster, the issue is still controversial. In effect, molecular phylogenetic studies performed to date generated ambiguous results since tree topologies depend on the kind of molecular marker employed. Curiously, even though mitochondrial DNA has become a popular marker in evolutionary biology and population genetics, none of the more than twenty *Drosophila* mitogenomes assembled so far belongs to this cluster. In this work we report the assembly of six complete mitogenomes of five species: *D. antonietae*, *D. borborema, D. buzzatii*, *D. seriema* and two strains of *D. koepferae*, with the aim to revisit the phylogenetic relationships and divergence times by means of a mitogenomic approach. The recovered topology using complete mitogenomes gives support to the hypothesis of the monophyly of that the *D. buzzatii* cluster and shows two main clades, one including *D. buzzatii* and *D. koepferae* (both strains) and the other the remaining species. These results are in agreement with previous reports based on a few mitochondrial and/or nuclear genes but in conflict with the results of a recent large-scale nuclear phylogeny, suggesting that nuclear and mitochondrial genomes depict different evolutionary histories.

## Introduction

Nowadays, almost every mitochondrial genome, called mitogenome, can be assembled directly from genome or even transcriptome sequencing datasets [1, 2]. The exponential development of next-generation sequencing (NGS) technologies, together with efficient bioinformatic tools for the analysis of genomic information make possible the fast and cheap assembly of mitochondrial genomes, giving rise to the emergence of the mitogenomics era [3]. Mitogenomics has been very useful in illuminating phylogenetic relationships at various depths of the Tree of Life, e.g. among early branching of metazoan phyla [4], among crocodilians and their survival in the Cretaceous-Tertiary boundary [5], Primates [6], the largest clade of freshwater actynopterigian fishes [7] and Anura, the largest living Amphibian group [8]. Also, mitogenomic approaches have been used to investigate evolutionary relationships in groups of closely related species (e.g. [9]). In animals, the mitochondrial genome has been a popular choice in phylogenetic and phylogeographic studies because of its mode of inheritance, rapid evolution and the fact that it does not recombine [10]. Such physical linkage implies that all regions of mitogenomes are expected to produce the same phylogeny. However, the use of different regions of the mitochondrial genome or even the complete mitogenome may lead to incongruent results [11], suggesting that mitogenomics sometimes may not reflect the true species history but rather the mitochondrial history [12–16]. Inconsistencies across markers may result from inaccurate reconstructions or from actual differences between genes and species trees. In fact, most methods do not take into consideration that different genomic regions may have different evolutionary histories, mainly due to the occurrence of incomplete lineage sorting and introgressive hybridization [17–19].

Since the last century, the *Drosophila* genus has been extensively studied because of the well-known advantages that several species offer as experimental models. A remarkable feature of this genus, that comprises more than two thousand species [20], is its diverse ecology: some species utilize fruits as breeding sites, others flowers, tree sap fluxes and cacti (reviewed in [21–24]). The adoption of decaying cacti as breeding sites occurred more than once in the evolutionary history of *Drosophilidae* [25, 26] and is considered a key innovation in the diversification and the invasion of American deserts by species of the *Drosophila repleta* group (*repleta* group from hereafter) [25]. Most species of this group are capable of developing in necrotic cactus tissues while feeding upon cactophilic yeasts associated to the decaying process [27–34].

The *repleta* group comprises more than one hundred species [22, 35-38], however, only one of the more than twenty complete (or nearly complete) *Drosophila* mitogenomes assembled so far belongs to a species of this group (checked in GenBank, March 28, 2019), *Drosophila mojavensis* (GenBank: BK006339.1). The latter, the first cactophilic fly to have a sequenced nuclear genome [39], is a member of the *D. mullerii* complex, an assemblage of species that belongs to the *D. mulleri* subgroup, one of the six species subgroups of the *repleta* group [36].

The *D. buzzatii* complex is the sister group of the *D. mulleri* complex [25]. It diversified in the Caribbean islands and South America, giving rise to the *D. buzzatii* (*buzzatii* cluster from hereafter)*, D. martensis* and *D. stalkeri* clusters [40]. The former is an ensemble of seven closely related species, *D. antonietae* [41], *D. borborema* [42], *D. buzzatii* [43], *D. gouveai* [41], *D. koepferae* [44], *D. serido* [42], and *D. seriema* [45]. All species are endemic to South America (Fig 1), except the semi-cosmopolitan *D. buzzatii* that reached a wide distribution following human mediated dispersion of prickly pears of the genus *Opuntia* (Caryophillales, Cactaceae) in historical times [34, 46, 47]. These species inhabit open areas of sub-Amazonian semidesertic and desertic regions of South America, where flies use necrotic cactus tissues as obligatory feeding and breeding resources [34, 48]. Regarding host plant utilization, *D. buzzatii* is an *Opuntia* specialist [30], a condition considered as ancestral [25]. However, *D. buzzatii* has also been recovered from necrotic columnar cacti [34]. The remaining species are mainly columnar dwellers though *D. antonietae*, *D. serido* and *D. koepferae* can also emerge marginally from rotting prickly pears [48].

**Fig 1.**
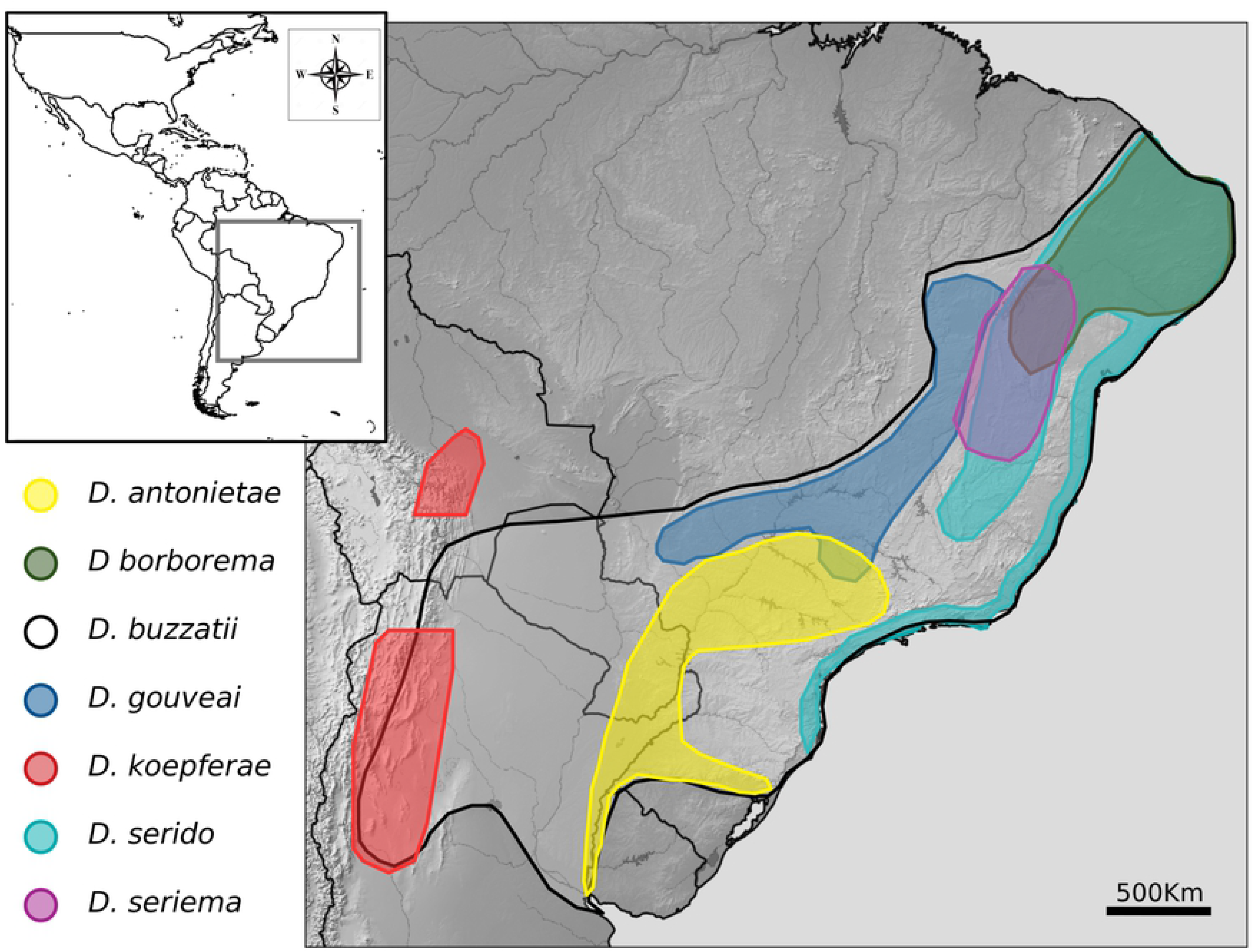
Geographical distribution of *buzzatii* cluster species. Modified from [49].

Species of the *buzzatii* cluster are almost indistinguishable by external morphology, however, differences in the morphology of male intromittent organ (aedeagus) and polytene chromosomes banding patterns provide clues to species identification (reviewed in [34, 47, 50]). The cluster has been divided into two groups based on aedeagus morphology, the first includes *D. buzzatii* and the remaining species compose the so-called *Drosophila serido* sibling set -*serido* sibling set from hereafter-[47]. In turn, the analysis of polytene chromosomes revealed four informative paracentric inversions that define four main lineages: inversion *5g* fixed in *D. buzzatii*, *2j^9^* in *D. koepferae*, *2x^7^* shared by *D. antonietae* and *D. serido*, and *2e^8^* shared by *D. borborema*, *D. gouveai*, and *D. seriema* [40, 51]. However, neither genital morphology nor chromosomal inversions are useful to discern the basal relationships within the cluster.

Pre-genomic phylogenetic studies based on a few molecular markers generated debate since different tree topologies were recovered depending on the molecular marker used. On one hand, the mitochondrial c*ytochrome oxidase I* (*COI*) and the *X-linked period* give support to the hypothesis of two main clades, one integrated by *D. buzzatii* and *D. koepferae* and another comprising the remaining species [47, 52, 53]. On the other hand, trees based on a few nuclear and mitochondrial markers support the hypothesis that *D. koepferae* is part of the *serido* sibling set [25, 54]. Moreover, a recent genomic level study using a large transcriptomic dataset supports the placement of *D. koepferae* as part of the *serido* sibling set and *D. buzzatii* as sister to this set [49]. However, phylogenetic relationships within the *serido* sibling set could not be ascertained despite the magnitude of the dataset employed by Hurtado and co-workers (2019). Thus, our aim is to shed light on the evolutionary relationships within the *buzzatii* by means of a mitogenomic approach.

In this paper, we report the assembly of the complete mitogenomes of *D. antonietae*, *D. borborema, D. buzzatii*, *D. seriema* and two strains of *D. koepferae*, together with the corresponding gene annotations. Here we also present a mitogenomic analysis that defines a different picture of the relationships within the *buzzatii* cluster with respect to the results generated with nuclear genomic data. Finally, we discuss possible causes of the discordance between nuclear and mitochondrial datasets.

## Material and Methods

### Species selection

The mitochondrial genomes of six isofemale lines of five species of the *buzzatii* cluster, for which NGS data are available, were assembled for the present study: *D. borborema* (obtained from Stock Center, derives from collections in Morro do Chapéu-Bahía State, Brazil)*, D. antonietae* (collected in Martín García Island, Argentina)*, D. buzzatii* (derives from collections in Spain by A. Ruiz); *D. seriema* (derived from collections in Serra do Cipó, Bahía State, Brazil) and two *D. koepferae* strains (*D. koepferae11* derives from collections in Bolivia by A. Fontdevila and A. Ruiz, and *D. koepferae7.1* collected in Vipos, Tucumán, Argentina by J. Hurtado and E. Hasson). The rationale of including these *D. koepferae* strains is motivated by previous protein electrophoresis work showing a certain degree of genetic divergence between Bolivian and Argentinian populations higher than between conspecific populations in other species [44]. In addition, we also included four species of the subgenus *Drosophila*, for which assembled mitogenomes are available, as outgroups in the phylogenetic analyses: *D. grimshawi* (GenBank: BK006341.1), *D. littoralis* (GenBank: NC_011596.1), *D. virilis* (GenBank: BK006340.1) and *D. mojavensis* (GenBank: BK006339.1).

### In silico mtDNA reads extraction

Whole genome sequencing (WGS) and RNA-seq data for *D. antonietae*, *D. borborema* and both strains of *D. koepferae* were generated in our laboratory ([33, 49], Moreyra & Hasson unpublished data), except for *D. seriema* and *D. buzzatii* for which mitochondrial reads were retrieved from the Genome sequencing of *D. seriema* deposited in Sequence Read Archive database (SRA accession ID: ERX2037878) [55] and the *D. buzzatii* genome project (https://dbuz.uab.cat), respectively. For each species mitochondrial reads were extracted from genomic and transcriptomic (when available) datasets. Bowtie2 version 2.2.6 [56] was first used with parameters by defaults (end-to-end sensitive mode) to map reads to the mitochondrial genome of *D. mojavensis*, the closest relative of *buzzatii* cluster species available, as reference. Next, only reads that correctly mapped to the reference genome were retained using Samtools version 1.8 [57]. Finally, mapped reads from genomic and transcriptomic datasets were combined to generate a set of only mitochondrial reads.

### Mitochondrial reference genome assembly

It is well known that at least 25% of NGS reads are of mitochondrial origin [3]. Therefore, after the mapping process it is possible to attain a coverage ranging from 2000x to more than 20000x for mitogenomes. In order to avoid miss-assemblies caused by the large number of reads, several coverage datasets were generated by random sampling. Then, a two-step assembly procedure was adopted for each coverage dataset based on recommendations of MITObim package version 1.8 [1]. In the first step, MIRA assembler [58] was run using the mitogenome of *D. mojavensis* as reference to build a new template from conserved regions. In the second step, the MITObim script was applied to the new template to reconstruct the entire mitochondrial genome by mapping the corresponding reads running a maximum of ten mapping iterations. All the different coverage assemblies were aligned with Clustalw2 version 2.1 [59] and, then, a consensus assembly was generated considering a sequence representation threshold of 60% not allowing gaps. This pipeline was employed for the assembly of the mitogenomes of all strains.

### PCR amplification, Sanger sequencing and consensus correction

Mitogenomes assembly coverage averaged more than 20000x, however, three regions including parts of *COI*, *NADH dehydrogenase subunit 6* (*ND6*) and *large ribosomal RNA* (*rRNAL)* genes, presented low read representation in all species, producing miss-assemblies and fragmentation. These regions were PCR-amplified with GO taq Colorless Master Mix by Promega using primers designed for regions conserved across the six mitogenomes assembled in this study (data in S1 Text). PCR amplifications included an initial denaturation at 94°C for 90 s, followed by 25 cycles of denaturation at 94°C for 45 s, annealing at 62°C for 50 s, extension at 72°C for 1 min and a final 4 min extension. PCR fragments were sequenced in both directions on an ABI-3130xl (Genetic Analyzer). Sequences were analyzed and filtered using Mega X software [60] and, finally, merged with the assemblies.

### Genome annotation and bioinformatic analyses

The six new assemblies were annotated in the MITOS web server (http://mitos.bioinf.uni-leipzig.de) [61], using the invertebrate mitochondrial genetic code and default parameter settings. The position and orientation of annotations were examined by mapping reads to mitogenomes with Bowtie2 [56] and visualization conducted with IGV version 2.4.10 [62]. In addition, nucleotide composition and codon usage were analyzed using MEGA version 10 [60]. A homemade python package (available upon request) was developed to compute estimates of pairwise nucleotide divergence (π) between *buzzatii* cluster species, and to visualize variation of π along the mitogenomic alignment of the cluster. Similar estimates were included for the *D. melanogaster* subgroup, a well-studied group of species, to compare divergence patterns along the mitochondrial DNA with the *buzzatii* cluster. To this end, mitogenomes of *D. melanogaster* (KJ947872.2), *D. erecta* (BK006335.1), *D. simulans* (NC_005781), *D. sechellia* (NC_005780) and *D. yakuba* (NC_001322.1) were aligned, and π estimates were obtained as described above. Synonymous (d_S_) and non-synonymous substitution rates (d_N_) were also estimated for each mitochondrial protein coding gene (PCG) using PAML 4.8 [63]. These estimates, as well as, the ω ratio (d_N_/d_S_) were obtained separately for both the *buzzatii* cluster and the *melanogaster* subgroup sequence alignments. Multiple sequence alignments of each coding gene were obtained with Clustalw2 version 2.1 [59].

### Phylogenetic analyses

Phylogenetic analyses were conducted considering PCGs, ribosomal genes (rRNAs), transfer RNA genes (tRNAs) and intergenic regions (excluding the control region) of the 6 mitogenomes plus the sequences of the outgroups *D. virilis*, *D. grimshawi*, *D. litoralis* and *D. mojavensis* (see details in species selection section). The aligment of the ten mitogenomes was performed with Clustalw2 version 2.1 [59]. The flanking sequences that correspond to the control region and portions of the alignment presenting abundant gaps were manually removed with Seaview version 4 [64]. The final alignment was used as input in PartitionFinder2 [65] to determine the best partition scheme and substitution models, considering separate loci and codon position (in PCGs), which were used in Bayesian Inference and Maximum Likelihood phylogenetic searches. In the Bayesian Inference approach, executed with MrBayes version 3.2.2 [66], both substitution model and parameter estimates were unlinked. Then, two independent Markov Chain Monte Carlo (MCMC) were run for 30 million generations with three samplings every 1000 generations, giving a total of 30,000 trees. Tracer version 1.7.1 [67] was used to assess the convergence of the chains mixing, where all parameters had ESS>200 (effective sample sizes), 25% of the trees were discarded as burn-in and the remaining trees were used to estimate a consensus tree and the posterior probability of each clade. The consensus tree was plotted and visualized with FigTree version 1.4.4 (https://github.com/rambaut/figtree/releases) [68]. Maximum Likelihood searches were performed in 2,000 independent runs using RAxML version 8.2.11 [69], applying the rapid hill climb algorithm and the GTR+GAMMA model, considering the partition scheme obtained with PartitionFinder2. Two thousand bootstrap replicates were run to obtain clade frequencies that were plotted onto the tree with highest likelihood. Tree and bootstrap values were visualized with Figtree version 1.4.4 [68]. Bayesian Inference searches for each PCG were individually done to seek for correlations with the topology recovered using the complete mitogenome. The GTR-GAMMA model, together with the same parameters and evaluation detailed before were applied on each MCMC.

### Divergence time estimation

Divergence times were estimated using the same methodology as in Hurtado et al., (2019). Four-fold degenerate third codon sites (putative neutral sites) of PCGs were extracted from the alignment and Bayesian Inference searches were run using BEAST version 1.10.4 [70]. A strict clock was set using a prior for the mutation rate of 6.2×10^−07^ per year (standard deviation of 1.89×10^−07^), as was empirically estimated for mitochondrial DNA in *Drosophila melanogaster* [71]. In addition, a birth-death process with incomplete sampling and a time of 11.3 myr (confidence interval ranging from ~9.34 to ~13) [25] to the root were defined as tree priors. Two MCMC were done in 30 million generations with tree sampling every 1000 generations. Tracer [67] was used to evaluate the convergence of the chains, discarding 10% of the total trees (burn-in). The information of the recovered trees was summarized in one tree applying LogCombiner and TreeAnnotator version 1.10.4 (available as part of the BEAST package), including the posterior probabilities of the branches, the age of the nodes, and the posterior estimates and HPD limits of the node heights. The target tree was visualized using FigTree [68]. Only *D. mojavensis* was included as outgroup in this analysis to minimize problems of among-taxa rate variation given by the large divergence between the *buzzatii* cluster and the rest of the species already sequenced, together with the lack of time point calibrations and accurate mutation rates.

## Results

### Mitogenomes characterization, nucleotide composition and codon usage

The length of the assembled mitogenomes varied from 14885 to 14899bp among the six strains reported in this paper. Mitogenomes consisted of a conserved set of 37 genes, including 13 PCGs, 22 tRNAs and 2 rRNAs genes, with order and orientation identical to *D. mojavensis*. Several short non-coding intergenic regions were also found. Twenty-three genes were found on the heavy strand (+) and fourteen on the light strand (-). Detailed statistics about metrics and composition of the mitogenomes are shown in Table 1.

**Table 1.** Composition of mitochondrial elements in the species assemblies of the *Drosophila buzzatii* cluster.

|  | Species assembly |  |  |  |  |  |
| --- | --- | --- | --- | --- | --- | --- |
| Statistics | *D. antonietae* | *D. borborema* | *D. buzzatii* | *D. koepferae11* | *D. koepferae7.1* | *D. seriema* |
| Total length | 14885 | 14889 | 14889 | 14892 | 14891 | 14891 |
| GC (%) | 23.36 | 23.22 | 23.60 | 23.20 | 23.20 | 23.26 |
| N's (%) | 0.00 | 0.00 | 0.00 | 0.00 | 0.00 | 0.00 |
| Intergenic (%) | 2.76 | 2.71 | 2.64 | 2.80 | 2.77 | 2.90 |
| tRNAs (%) | 9.82 | 9.81 | 9.80 | 9.81 | 9.81 | 9.80 |
| PCGs (%) | 72.98 | 73.04 | 73.15 | 72.95 | 72.97 | 72.85 |
| rRNAs (%) | 14.44 | 14.45 | 14.41 | 14.44 | 14.44 | 14.44 |

Overall nucleotide composition in PCGs ranged between 37.6-37.8% A, 37.2-37.9% T, 10.2-10.4% G, and 14.1-14.7% C. The thirteen PCGs were AT-biased as in the entire mitogenome, and the codon usage bias in each gene was greater than 0.50. The most frequently used codons were UUA (Leu), AUU (Ile), and UUU (Phe) in all cases. Codon usage information for each species is shown in Table in S1 Table.

### Genetic diversity among mitogenomes

Pairwise nucleotide diversity (π) estimates for both the *buzzatii* cluster and the *D. melanogaster* subgroup are shown in Fig 2. Though π values along the mitochondrial genome alignment were, on average, larger in the *melanogaster* subgroup than in the *buzzatii* cluster, pattern of variation of π along the entire molecules were very similar. Large and small ribosomal subunits (rRNAs) exhibited lower divergence values than the remaining mitochondrial genes in both groups. A substantial difference was found in the region encompassing *COIII*, *tRNA-G* and *ND3* genes. At these positions, from 5000 to 6000, nucleotide diversity was the highest in the *melanogaster* subgroup, showing an apparent increase represented by two high peaks absent in the *buzzatii* cluster. Considering genetic divergence within the *buzzatii* cluster (Fig 3), the lowest value of average pairwise nucleotide divergence was observed for the pair *D. borborema* and *D. seriema* (π = 1.91×10^−03^), while between *D. seriema* and *D. buzzatii* divergence was an order of magnitude larger (π = 2.73×10^−02^). Divergence between *D. koepferae* strains was surprisingly high (7.14×10^−03^). The complete set of divergence estimates in the *buzzatii* cluster is reported in Table in S2 Table. Substitution rates in synonymous and non-synonymous sites are listed in Table 2. The ratio d_N_/d_S_ (ω) varied from 0.003 to 0.060 among PCGs in the *buzzatii* cluster. The range in the *melanogaster* subgroup was similar, but with a lower upper bound (0.003 - 0.018). Two loci appear as outliers in the *buzzatii* cluster (*ATP8* and *ND2*), which apart from these two loci, has lower divergence values than in the *melanogaster* subgroup. In any case, the results suggest that purifying selection imposes strong constraints in the evolution of mitochondrial genes. Nonsynonymous (d_N_), synonymous (d_S_) and the ω ratios varied among PCGs (Table 2).

**Fig 2.**
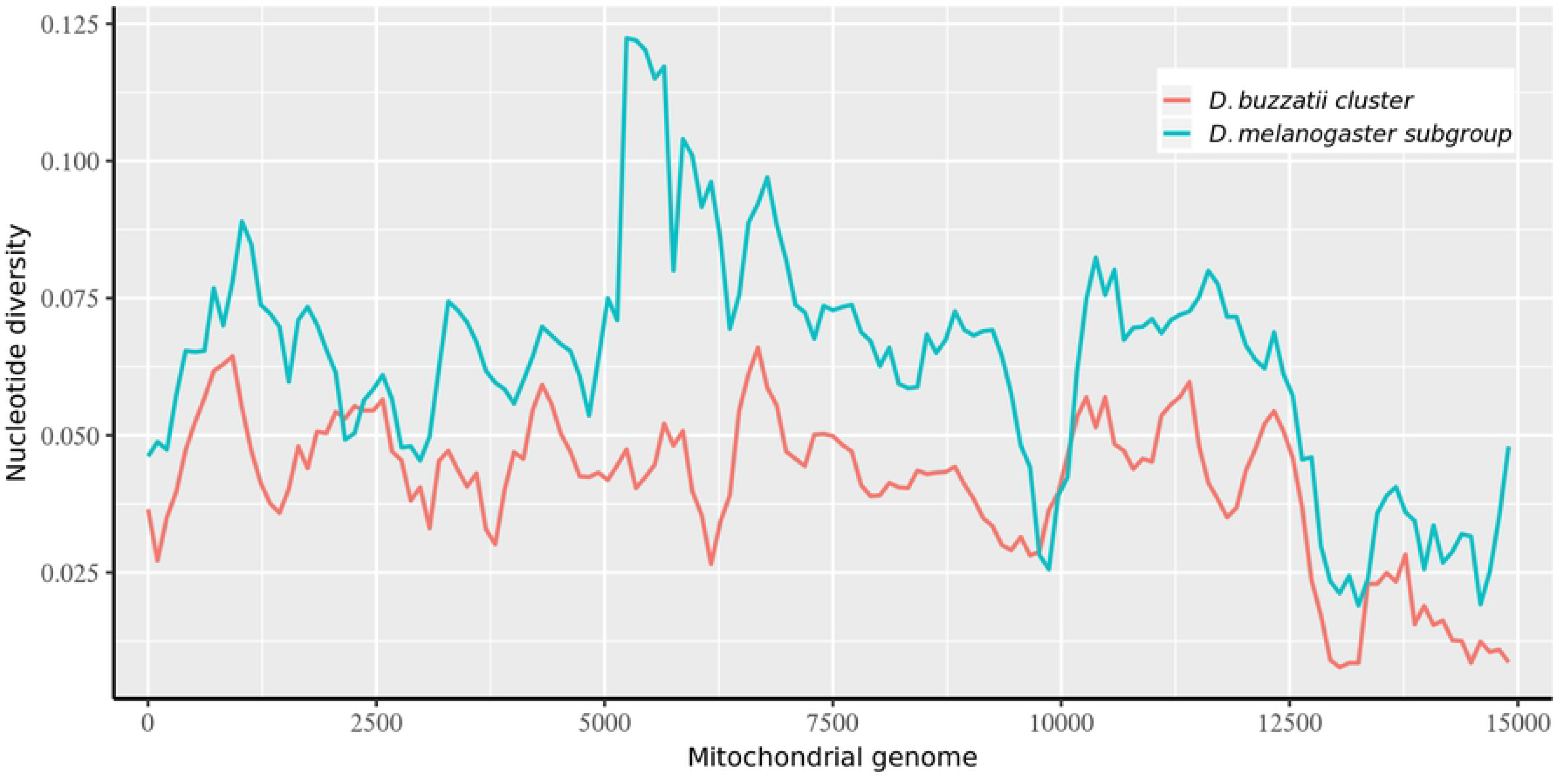
Nucleotide diversity (π) variation along the mitogenome, estimated for a sliding window of 500bp with an overlap of 100bp. π values for species belonging the *buzzatii* cluster and the *melanogaster* subgroup are represented independently.

**Fig 3.**
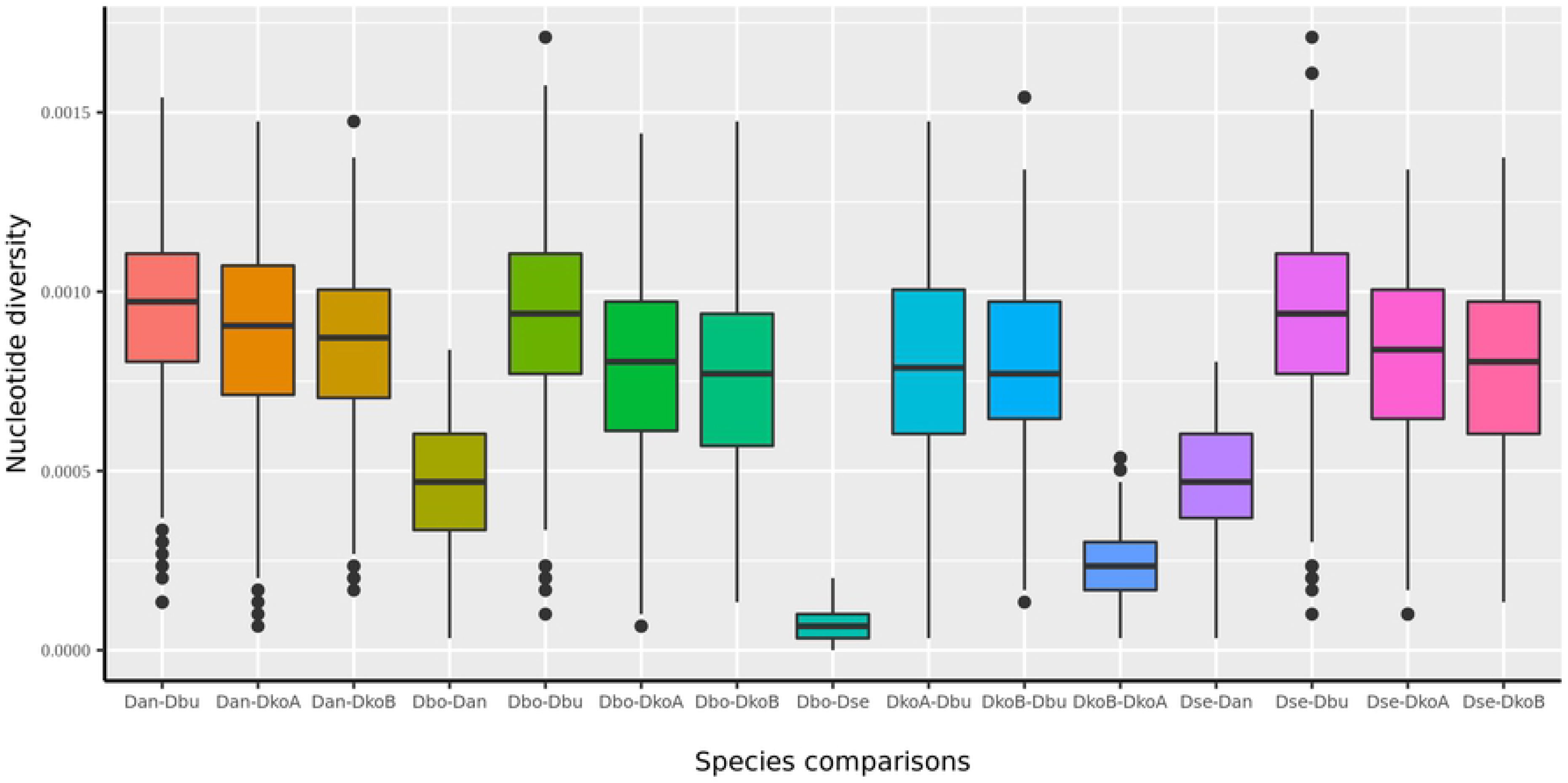
Pairwise comparison of nucleotide diversity between species belonging the *buzzatii* cluster.

**Table 2.** Estimates of non-synonymous (d_N_) and synonymous (d_S_) substitutions and their ratio (ω) among species of the *buzzatii* cluster and the *melanogaster* subgroup.

| PCG | *buzzatii* cluster |  |  | *melanogaster* subgroup |  |  |
| --- | --- | --- | --- | --- | --- | --- |
| | $\omega$ | $d_S$ | $d_N$ | $\omega$ | $d_S$ | $d_N$ |
| *ATP6* | 0.005 | 1.390 | 0.007 | 0.014 | 2.364 | 0.034 |
| *ATP8* | 0.060 | 0.506 | 0.031 | 0.014 | 4.892 | 0.071 |
| *CytB* | 0.005 | 1.463 | 0.008 | 0.010 | 2.556 | 0.027 |
| *COI* | 0.003 | 1.211 | 0.003 | 0.003 | 2.383 | 0.006 |
| *COII* | 0.005 | 1.058 | 0.005 | 0.008 | 2.295 | 0.018 |
| *COIII* | 0.006 | 1.277 | 0.008 | 0.012 | 1.975 | 0.024 |
| *ND1* | 0.003 | 4.332 | 0.012 | 0.006 | 4.490 | 0.028 |
| *ND2* | 0.036 | 1.102 | 0.040 | 0.016 | 2.368 | 0.039 |
| **ND3** | 0.009 | 3.657 | 0.034 | 0.016 | 3.085 | 0.051 |
| **ND4l** | 0.008 | 1.158 | 0.009 | 0.001 | 8.026 | 0.013 |
| **ND4** | 0.011 | 1.253 | 0.013 | 0.009 | 4.306 | 0.040 |
| **ND5** | 0.007 | 2.564 | 0.018 | 0.013 | 4.764 | 0.063 |
| **ND6** | 0.012 | 2.231 | 0.027 | 0.018 | 4.534 | 0.082 |

### Phylogenetics analyses

The sequences of the 13 PCGs, 22 tRNAs genes, 2 rRNAs genes, and intergenic regions were included in the alignment. Total length of the final matrix encompassing the ten mitogenomes was 15044 characters, from which 1950 were informative sites, 11583 conserved, and 1422 were singletons. Both Maximum Likelihood and Bayesian Inference phylogenetic analyses recovered the same highly supported topology that confirms the monophyly of the *buzzatii* cluster (Fig 4). Two main clades can be observed in the tree, one including both *D. koepferae* strains as sister to *D. buzzatii*, and the second, comprising *D. antonietae* as sister species of the sub-clade formed by *D. borborema* and *D. seriema*. The species selected as outgroups were allocated as expected, with *D. mojavensis* as the closest relative of the *buzzatii* cluster. We also performed a gene tree analysis using all PCGs (S1 Fig). We could only obtained trees for 7 genes out of the thirteen PCGs, given the lack of informative sites in the alignments of *ATP8*, *ATP6*, *ND3*, *ND4l*, *COII* and *COIII*. Only two (*ND1* and *ND5*) out of the seven recovered gene trees that showed the same topology as the complete mitogenome, while the remaining genes produced three (different) topologies. Trees obtained with *CytB* and *ND4* allocated *D. buzzatii* as sister of the *serido* sibling set which included *D. koepferae*. *COI* and *ND2* retrieved trees where *D. buzzatii* and *D. koepferae* exchanged positions in the tree, placing *D. koepferae* as the species closest to the putative ancestor of the cluster. *ND6* recovered two clades where *D. antonietae* was the sister of *D. buzzatii* and *D. koepferae* (both strains) in one clade, and the pair *D. borborema-D. seriema* composed the other.

**Fig 4.**
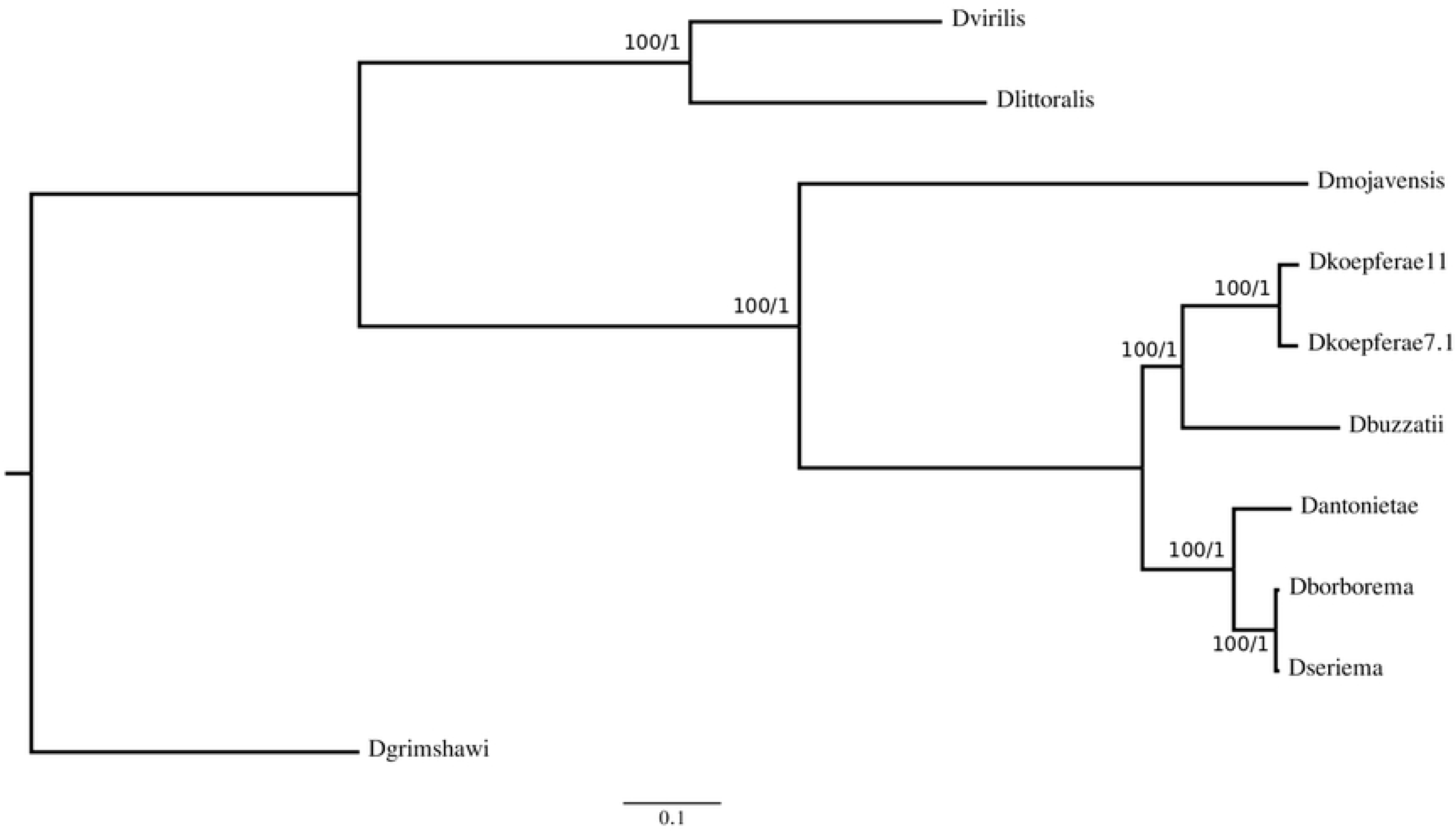
Phylogenetics hypotheses for the *buzzatii* cluster based on the entire sequence of the mitogenome (control region not included). Tree topology recovered by both Maximum Likelihood and Bayesian Inference searches. On each node is represented the bootstrap and the posterior probability, respectively.

### Divergence Times

PCGs contained 1201 4-fold degenerate sites in the mitogenomes of the *buzzatii* cluster strains assembled in this study. The tree obtained in the divergence time estimation analysis (Fig 5) was topologically identical to the trees obtained in the phylogenetic analyses using complete mitogenomes (see Fig 4). Divergence time estimations showed that the *buzzatii* cluster diverged in the Early Pleistocene, 2.11 Myr ago, and the split with the *D. mojavensis* common ancestor occurred 10.63 Myr ago in the Miocene. Our results also indicated that the clade containing *D. antonietae, D borborema* and *D. seriema*, is younger than the clade composed by *D. buzzatii* and *D. koepferae*. In addition, the split between *D. borborema* and *D. seriema* is quite recent, about ~50000 years ago, in the Late Pleistocene, even more recent than the split of *D. koepferae* strains that diverged ~310000 years ago, in the Middle Pleistocene.

**Fig 5.**
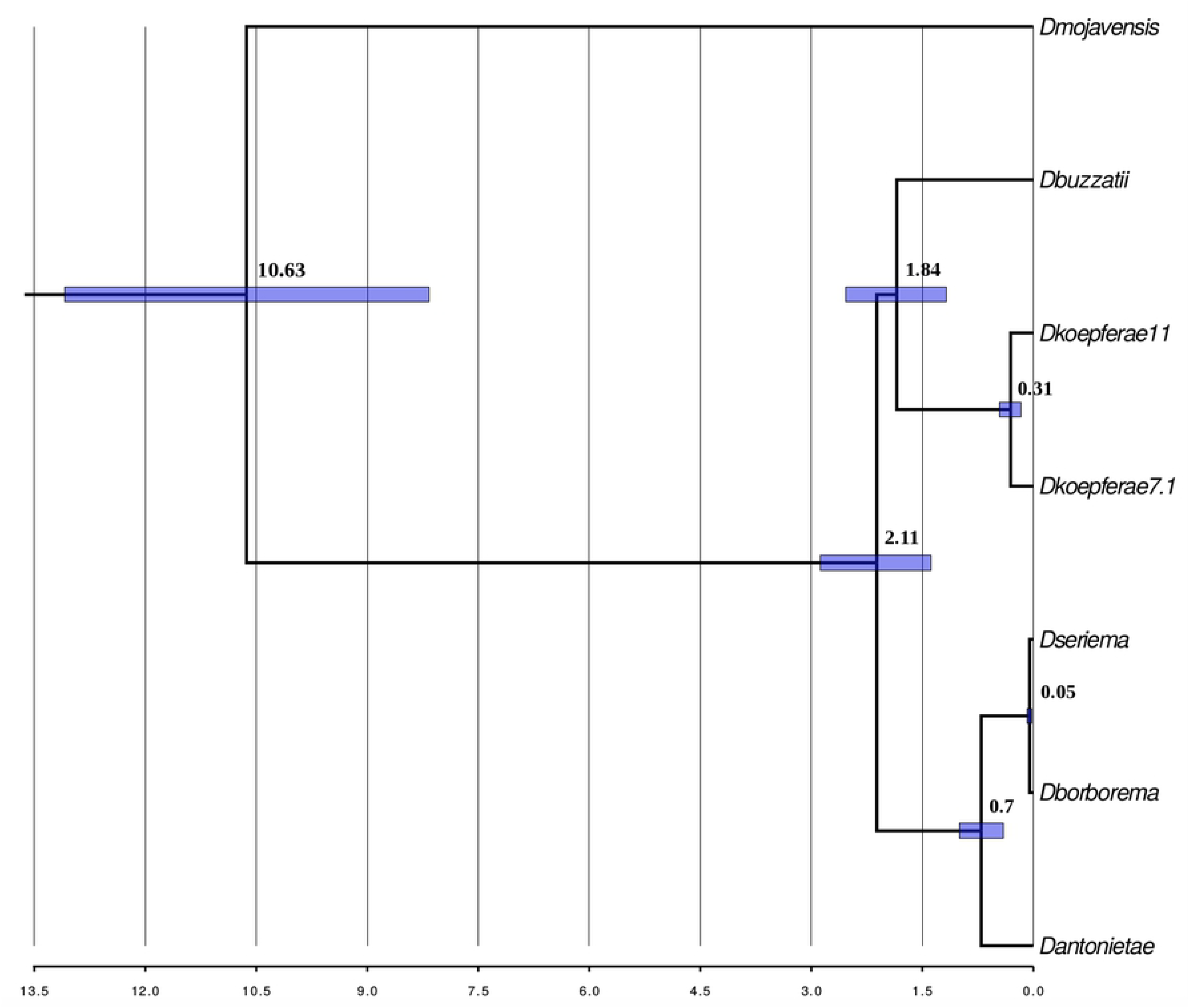
Divergence times for the *buzzatii* cluster drawn on a Bayesian Inference tree. Numbers on each node are the time estimates. Blue bars represent the 95% confidence intervals of estimates.

## Discussion

In this paper, we report six newly assembled mitochondrial genomes of five cactophilic species of the *buzzatii* cluster. Our aim is to revisit the phylogenetic relationships by means of a mitogenomic approach in this set of closely related species in active cladogenesis.

Structural analyses showed that the newly assembled mitogenomes share molecular features with animal mitochondrial genomes sequenced so far [72]. All assembled mitogenomes contain the same set of genes usually found in animal mitochondrial genomes. Gene order and orientation, as well as the distribution of genes on the heavy and light strands are identical to the mitogenome of the closest relative *D. mojavensis* and other drosophilids [9]. Analysis of overall nucleotide composition of mitogenomes and PCGs revealed the typical AT-bias found in *Drosophila* mitogenomes. Codon usage is highly biased suggesting that synonymous sites cannot be considered strictly neutral and that some sort of natural selection for translational accuracy governs codon usage [73].

Phylogenetic analyses based either on complete mitogenomes or four-fold degenerate sites (for divergence time estimations), retrieved a high-confidence tree, suggesting that the cluster is composed by two main clades, one including *D. buzzatii* and *D. koepferae* (both strains) and another comprising *D. antonietae*, *D. seriema* and *D. borborema*. These results are consistent with previous work based on single mitochondrial genes [52, 74], but inconsistent with phylogenetic studies based on both a small set of nuclear and mitochondrial genes [25] and a large set of nuclear genes-see below-[49]. Interestingly, the topology showing these two clades was only recovered in two (*ND1* and *ND5*) out of seven trees based on individual PCGs, whereas the remaining trees produced either a novel topology or a topology consistent with the phylogeny reported in Hurtado et al., (2019).

The lack of recombination causes mitochondrial DNA to be inherited as a unit, thus, trees obtained with individual mitochondrial genes are expected to share the same topology and to be consistent with the trees obtained with complete mitogenomes. Thus, our results suggest that using individual genes not only produce different topologies but also a poor resolution of phylogenetic relationships. Such inconsistencies between complete mitogenomes and gene trees in phylogenetic estimation may result from inaccurate reconstruction or from real differences among gene trees. The first possible explanation is the simple fact that numbers of informative sites within a single locus are not enough to accurately estimate phylogenetic relationships, particularly in groups of recently diverged species (overall support for gene trees was poorer than for the tree based on complete mitogenomes). Secondly, heterogeneity in evolutionary rates among genes and/or differences in selective constraints along the mitogenome can also account for the inconsistencies [75–77]. As a matter of fact, we detected substantial variation of synonymous and nonsynonymous rates as well as of the ω ratio across PCGs. In addition, variation among oxidative phosphorylation complexes in the *buzzatii* cluster was high. The *ND* complex was, on average, less constrained than the *ATP* complex, *cytochrome b (CytB)* and *cytochrome oxidase* complex *(COI,COII & COIII)*, consistent with results reported in the *melanogaster* subgroup [9, 78]. Another factor that may lead to biased tree construction, particularly relevant for mitochondrial genes characterized by high substitution rates, is substitutional saturation [79]. A priori, saturation should not be problematic in recently diverged species, like the *buzzatii* cluster, however, saturation may be problematic in the estimation of divergence relative to the outgroup and, thus, for phylogenetic inference. The closest outgroup to the *buzzatii* cluster employed in our study is the *mulleri* complex species *D. mojavensis.*. Available evidence suggest that these complexes diverged ~10 MYA (but see a more recent estimate by Hurtado et al −2019-of 5.5Myr) suggesting that substitution saturation may lead to inaccurate phylogenetic reconstruction.

In this context, a recent report investigating the effect of using individual genes, subsets of genes, complete mitogenomes and different partitioning schemes on tree topology suggested a framework to interpret the results of mitogenomic phylogenetic studies [11]. The authors concluded that trees obtained with complete mitogenomes reach the highest phylogenetic performance and reliability than single genes or subsets of genes. Therefore, we consider that phylogenetic relationships inferred from complete mitogenomes reflect the evolutionary history of, at least, mitogenomes.

The phylogenetic relationships depicted by our mitogenomic approach are incongruent with a recent study based on transcriptomic data [49]. Based on a concatenated matrix of 813 kb uncovering 761 gene regions, the authors obtained a well-supported topology in which *D. koepferae* appears phylogenetically closer to *D. antonietae* and *D. borborema* than to *D. buzzatii*, placing *D. buzzatii* alone as sister to the rest of the cluster. This topology is in agreement with male genital morphology, cytological and molecular phylogenetic evidence [25, 34, 54]. Nevertheless, the pattern of cladogenesis of the trio *D. koepferae-D. borborema-D. antonietae* could not be fully elucidated since a nuclear gene tree analysis yielded ambiguous results. As a matter of fact, the analysis of the 761 gene trees reported showed that about one third of the genes supported each one of the three possible topologies for the trio *D. koepferae-D. antonietae-D. borborema* indicating a hard polytomy [49]. In contrast, the early separation of *D. buzzatii* from the *serido* sibling set is supported by 97% of the genes and, surprisingly, none of the gene trees recovered the clade including *D. buzzatii* and *D. koepferae* as the sister group of the clade involving *D. antonietae* and *D. borborema* (J. Hurtado, F. Cunha-Almeida, E. Hasson, unpublished results) as suggested by the present mitogenomic approach.

Such mitonuclear discordance, has been reported in several animal species. A recent review lists several examples in animals [80]. Likewise, the literature in this respect is abundant in the genus *Drosophila*. Well-known cases are *D. pseudoobscura* and *D. persimilis* [81]; *D. santomea* and *D. yakuba* [82]; and *D. simulans* and *D. mauritiana* [83]. Mitonuclear discordance may be caused by incomplete lineage sorting (ILS) and/or introgressive hybridization. These two factors do not affect equally mitochondrial and nuclear genomes, ILS is more likely for nuclear genes, especially when the ancestral effective population size of recently diverged species was large [84, 85], while introgressive hybridization is expected to be prevalent in mitochondrial genomes given its lower effective population size [86]. If we accept that the topology based on nuclear genes is representative of the species-history (see also [48]), the closer similarity between *D. buzzatii* and *D. koepferae* mitogenomes is suggestive of gene flow between these largely sympatric species [34]. Thus, we suggest that *D. buzzatii* and *D. koepferae* lineages initially separated but then exchanged genes via fertile F1 females (males were likely sterile as expected by the ubiquitous Haldane rule) before finally separating less than 1.5 Myr ago. Not only the more recent mitogenomic ancestry is suggestive of gene exchange, also traces of introgressive hybriodization can still be detected in nuclear genomes [49].

In fact, phylogenetic, population genetic and experimental hybridization studies suggest a significant role of introgression in the evolutionary history of the *buzzatii* cluster. Phylogeographic studies revealed discordances between mitochondrial markers and genital morphology in areas of sympatry between species [52]. Likewise, interspecific gene flow has been invoked to account for shared nucleotide polymorphisms in nuclear genes in *D. buzzatii* and *D. koepferae* that cannot be accounted by ILS [87, 88]. Moreover, experimental hybridization studies have shown that several species of the *buzzatii* cluster can be successfully crossed, producing fertile hybrid females that can be backcrossed to both parental species. Interestingly, *D. koepferae* can be crossed with *D. antonietae*, *D. borborema*, *D. buzzatii* and *D. serido* [44, 47, 89-93].

Our estimates of divergence time are in conflict with previous studies. In general, previous estimates, based on individual or a few genes (either mitochondrial or nuclear) suggested an older origin of the cluster and deeper splitting times within the cluster when compared to the estimates based on transcriptomes and mitogenomes. In efect, Gómez & Hasson (2003) and Oliveira et al., (2012) dated the split of *D. buzzatii* from the remaining species of the cluster in ~4 or 4.6 Myr, respectively, whereas Manfrin et al.,’s (2001) estimates are even older, from 3 to 12 Myr for the most recent to the more ancient split. In contrast, putting apart the divergence time of the clade *D. buzzatii-D. koepferae* for the reasons discussed above, the radiation of the remaining three species seems to be extremely recent, less than 1 Myr ago (Fig 5) using mitochondrial genomes, which are similar to estimates based on transcriptomes [49]. However, it is worth mentioning that divergence times estimated in the present paper and by Hurtado et al. (2019), may be biased downwards since both are based on empirical mutation rates for nuclear and mitochondrial genes, respectively, calculated over 200 generations for *D. melanogaster* [71]. Thus, these results should be interpreted with caution in the light of evidence suggesting not only the time-dependence of molecular evolutionary rates but also that mutation rates obtained using pedigrees and laboratory mutation-accumulation lines, often exceed long-term substitution rates by an order of magnitude or more [77]. In this sense, an alternative method without a mutation rate prior, measured a rate four times lower that the reported in Haag-Liautard et al., (2008), consequently, yielding older divergence times (results not shown).

Even though divergence times estimates obtained in this study cannot be entirely compared to assessments based on nuclear genomic data and individual nuclear genes, given uncertainty of tree topology, they concur in the fact that species of the *buzzatii* cluster emerged during the Late Pleistocene in association with Quaternary climate fluctuations [48, 49, 74]. Moreover, in view of the obligate ecological association between *buzzatii* cluster species and cacti, the so-called Pleistocene “refuge hypothesis” is a suitable explanation for the diversification in this group in active cladogenesis. This hypothesis argues that Pleistocene glacial cycles successively generated isolated patches of similar habitats across which populations may have diverged into species [94, 95].

Available paleo-climatic evidence, consistent with the Pleistocene “refuge hypothesis”, can also account for the relatively deep intraspecific divergence between Bolivian and Argentinian *D. koepferae* strains. In effect, because Quaternary topographical patterns in the Central Andes have remained unchanged in the last 2-3 Myr, a plausible explanation for this late Pleistocene vicariant event is related with glacial-interglacial cycles [96]. Although the validity of the Pleistocene “refuge hypothesis” is controversial (cf. [97]) and few studies addressed specific hypotheses on how the Quaternary glacial-interglacial cycles impacted species diversification [98], our divergence time estimates between Bolivian and Argentinian *D. koepferae* suggest a role of climatic oscillations as a factor of ecogeographical isolation in the Central Andes during the Pleistocene. Moreover, paleo-climatological evidence suggest that the area inhabited by *D. koepferae* has been exposed to substantial climatic variations on timescales of 10^3^ to 10^5^ years related with Glacial-interglacial cycles. Thus, Andean north-south exchanges may have been alternately favored or disfavored by these Quaternary climatic oscillations. In fact, the estimated age of the vicariant event between the *D. koepferae* strains is tantalizingly coincident with the coldest phase of the Marine Isotopic Stage (MIS) 10, which corresponds to a glacial period that ended about 337,000 years ago [99]. The coldest period of the MIS 10 (recorded in global air and sea surface temperature and also the lowest atmospheric CO_2_ levels) occurred at 355.000 years, well within the confidence interval of our divergence time estimated between *D. koepferae* strains. In a global scale, glacial periods are primarily reflected in a lowering of air temperature but also in altered patterns of precipitation in the both sides of the Central Andes [100] which were in turn the main drivers of vegetation changes [101] including the appearance of South American columnar cacti [102]. Besides the impact on air temperature, periods of ice advance in the Central Andes generally were periods of negative water balance in the Pacific coastal regions west to the Central Andes [103], and a positive water balance in the Central Andes, as evidenced by deeper and fresher conditions in Lake Titicaca [104] (see S2 Fig). Thus, during the colder and wetter phases of the MIS 10 in the Central Andes, species distributions may had suffered a general contraction towards the southern and northern lowland warmer refugia between 1000-2000 m, whereas a general worsening condition occurred in higher western elevations. North and south refugia were probably separated by a gap of low suitability represented by the steep gradient of the eastern flank of Eastern Andes between 22-24°S, which represents today a region of strong W-E precipitation gradient. The MIS 10 glacial cycle has a particular structure since it does not have a pronounced interstadial (relative warmer) conditions in the mid-cycle [105], providing a prolonged, effective “soft” dispersal barrier that affected the distribution of *D. koepferae*.

Finally, our present study indicates the need of counting with the mitogenomes of the other Brazilian species *D. gouveai and D. serido* to achieve a deeper understanding of the evolutionary history of the cluster. A comparative analysis including the complete mitogenomes of all species may help to disentangle the intricate relationships in the *buzzatii* cluster.

## Acknowledgements

Nicolás N. Moreyra is recipient of a PhD scholarship awarded by CONICET and ANPCyT. JM, JH, FA and EH are fellows of CONICET.

## Supporting Information

### Captions

**S1 Text.** Pair of primers designed for regions conserved across the six mitogenomes.

**S1 Table.** Codon usage for each mitogenome of the *buzzatii* cluster species.

**S2 Table. Genetic divergence among species of the *buzzatii* cluster.** Estimates are shown for each pairwise comparison between species.

**S1 Fig.** Phylogenetic hypotheses for the *buzzatii* cluster species recovered by each mitochondrial gene using Bayesian Inference searches.

**S2 Fig. Paleoclimatic records of the last 500,000 years.** Ages in the top are indicated as 10^3^ years (kyrs). Gradated shading area indicates divergence age estimates. Marine Isotope Stages (MIS) are labeled according to Lisiecki and Raymo (2005). Shaded vertical areas correspond to glacial periods whereas white areas correspond to interglacials or interstadials. Glacial periods correspond to cold and dry conditions in the western slopes of the Western Andes, and cold and wetter conditions in the eastern slopes of the Eastern Andes and the Altiplano. **A**. Globally-averaged surface air temperature anomaly reconstructed from proxy and model data for the last eight glacial cycles [106].

**B.** CO2 concentration based on Vostok Ice Core data [107]. **C.** Iron accumulation rates (AR Fe) reflecting changes in terrigenous sediment input to ODP Site 1239D, Equatorial Pacific [108]. **D.** % of CaCO3from Site LT01-2B indicating changes in water balance at Lake Titicaca Basin, Bolivia (modified from 104).

